# Replacing fish meal with untreated and enzymatically treated torula yeast (*Cyberlindnera jadinii*) affects pelleting die flow resistance and physical properties of the feed pellets

**DOI:** 10.1101/2025.02.21.639432

**Authors:** Dejan Dragan Miladinovic, Carlos Salas-Bringas, Esther Julius Mbuto, Pashupati Suwal, Odd Ivar Lekang

## Abstract

Among novel feed ingredients, yeast has gained significant importance. Fishmeal typically provides superior feed conversion ratios (FCR) in farmed aquatic organisms compared to most alternative ingredients. However, published data suggests that the yeast *Cyberlindnera jadinii* is a promising alternative. This study demonstrates that replacing fishmeal with torula yeast, either untreated or treated with protease and endo-exo 1.3-β-glucanase, reduces feed material resistance in the pelleting die during pellet discharge. It also investigates how enzyme-treated yeast affects pellet quality. Experiment 1 examined changes in flow resistance and pellet quality when fishmeal was replaced by yeast. Experiment 2 assessed the impact of enzyme-treated yeast on pellet flow resistance and physical quality. Results showed a significant increase in flow resistance in diets with 20% yeast and enzyme-treated yeast. The tensile strength of pellets with 20% yeast increased in both experiments. Pellets with 10%, 20%, and 100% yeast exhibited aquaphobic behavior compared to those without yeast, which benefits feed consumption, reduces wastage, and supports sustainable feed manufacturing. However, pellets with 10% and 20% enzyme-treated yeast showed lipophobic behavior, which is less ideal during post-production processes. Enzymatic treatment reduced underwater swelling of pellets with 10% and 20% yeast. Minimal enzymatic hydrolysis is recommended to lower underwater disintegration. The longitudinal surface roughness of pellets decreased with enzyme-treated yeast, with pellets containing 20% treated yeast showing the lowest surface roughness.

## 1. Introduction

Expansion of aquaculture requires increased quantities of alternative and sustainable protein sources (Aslaksen *et al*., 2007). Fishmeal remains an important part of aquatic-feed diets and the feed manufacturing industry is a substantial consumer of fishmeal (Tacon and Metian, 2015; Malcorps *et al*., 2019). Therefore, this growing industry, together with research institutions, tries to target the best possible alternative candidates to replace fishmeal (Froehlich *et al*., 2018; Pelletier *et al*., 2018). In the last decade, various yeast species gained attention as a good source of nutritionally valuable raw materials for aquatic feeds (Langeland *et al*. 2014; Vidakovic *et al*., 2015; Huyben *et al*. 2017; Vidakovic *et al*., 2019; Agboola *et al*. 2020). Yeasts have high nutritional value based on protein with an optimal composition of amino acids for farmed aquatic animals such as tilapia (Olvera Novoa *et al*. 2002), salmon (Øverland *et al*. 2013) and shrimp (Gamboa-Delgado *et al*. 2015). Yeasts can be successfully used as alternatives to fishmeal or soybean meal up to 24% without impacting the growth (Guo *et al*., 2019) or health condition (Jin *et al*., 2018) of the aquatic animals. It is common knowledge that replacing conventional feed ingredients in complete feed diets can influence the physical quality loss of the feed. Loss of feed nutrients in the water may contribute to water pollution (Tantikitti, 2014) and consequently increase stress on the farmed aquatic animals and reduce production (Ferreira *et al*., 2011).

Generally, the acceptable physical properties of feed pellets are indicators of improved feeding efficiency and convenient feed handling. Such evaluation may be done by characterizing the influence of added novel material in the feed matrix on durability, hardness, flow resistance of the material in the pelleting die during pelleting, hydrophilicity, and pellet swelling underwater. Evaluating the surface characteristics and hygroscopicity of the feed pellets containing a selected alternative ingredient may help determine the functionality of that ingredient. This would further contribute to better feed manufacturing, better shelf-life estimation, texture evaluation of the feed product, and quality control (Stokes *et al*., 2013).

During the compaction of the feed mash, the resistance to the flow of the raw materials can influence the consumption of electrical energy, consistency, and physical quality of the final product. When feed material resists flow during processing this correlates with the temperature change in the die. Heating in the pelleting die together with sufficient water content in the feed mash can change the nutrients during processing (Hoseney *et al*., 1992). Controlled processing variables, such as the shearing of the material in the pelleting die, can alter the microstructure of the final product (Hermansson, 2000). Shearing during the pelleting process significantly impacts the microstructure of the pellets by breaking down larger particles and redistributing them more uniformly throughout the compacting matrix. This mechanical action reduces the size of air pockets and enhances the density and cohesion of the pellets. As a result, the pellets exhibit improved physical properties such as increased strength and durability. These changes are crucial for the quality of the pellets, as a more uniform and denser microstructure ensures better handling and reduced fines. Also, enzymatic treatment during feed manufacturing has a significant ability to better use feed ingredients for improved nutritional and physical quality of feed (Bedford and Partridge, 2011). When yeast is included in the feed, the components of the cell wall, such as β-glucans and mannoproteins, can absorb water and can swell. This absorption leads to the formation of a gel-like matrix within the feed mass which can form viscous environment. As these components interact with other particles, they can create a more cohesive and viscous network (Feldmann, 2012). The chemical composition of the yeast cell wall is mostly consisted of mannoproteins, from 30% to 50%; β-(1,6)-glucan from 5% to 10%; β-(1,3)-glucan from 30% to 55% and up to 2% chitin of cell-wall dry weight (Klis *et al*. 2002; Kurtzman *et al*., 2011). Changing the structure of the yeast cell wall with enzymes and their hydrolytic activity could reduce viscosity during pelleting. Structural change of the yeast cell wall can occur when there is increased hydrolytic influence with the enzymes protease and beta-glucanase on dominating molecules such as protein and β-(1,3)-glucan. When the chemical bonds in the mannoproteins and β-(1,3)-glucan molecule are destroyed, the viscosity decreases (Li *et al*. 2025), and hence the packing of the feed material during pelleting may change. This can lead further to changes of the physical properties of the feed pellets. However, short-time conditioning during feed pelleting with low water content would potentially be limiting factors for making any change in the chemical or physical aspects connected to the feed quality. There is a lack of published data on how adding different enzymes relates to the flow resistance of the pellets through the pelleting die and the physical quality of the feed pellets, and a lack of reported evidence on how the enzymes exo-endo-1,3-β-glucanase and protease affect the compacted feed material during pelleting.

If the enzymes in a low water content environment are modifying the biochemical properties of the feed ingredients, this could lead to an altered flow of the material during compaction and thus modify the physical properties of the feed pellets (Karan et al., 2012).

If the flowability changes due to the changed pressure at incipient flow (p_max_) in the pelleting die, the pelleting equipment’s power consumption also changes. This leads to a change in the physical properties of feed pellets (Miladinovic *et al*. 2021). The entire spectra of different analyses may contribute to the overall understanding of the physical quality of pellets. Water activity (a_w_) is a quality parameter influencing final physical and nutritional feed quality (Lowe and Kershaw, 1995). If the water activity is too high, it can lead to moisture migration within the feed, causing undesirable textural changes. This can result in caking of the feed, making it less palatable and harder to handle. Physical characteristics of the feed at a given a_w_ may be affected by the thermodynamics of a feed ingredient (Slade and Levin, 1991). Pressure at incipient flow (p_max_) during pelleting is influenced by water activity, as higher water activity can reduce the material’s resistance to flow, making it easier to compress and form pellets. These factors are crucial because they affect the pelleting process’s efficiency, the pellets’ quality, and the energy consumption during production (Miladinovic *et al*. 2021).

This study investigates how replacing fish meal with the yeast *Cyberlindnera jadinii* influences the flow of feed mash in the pelleting die, with or without enzyme addition. It hypothesizes that using yeast and enzymes (protease and exo-endo 1-3-beta glucanase) can lower p_max_ during pelleting, potentially modifying the physical characteristics of feed pellets to enhance their durability and underwater stability, thereby improving their availability to farmed aquatic animals. This study comprised two experiments utilizing single-die pelleting to simulate commercial production conditions. The first experiment aimed to determine the optimal dosage of *Cyberlindnera jadinii* to enhance the physical quality of the pellets. The second experiment assessed the impact of replacing fishmeal with *Cyberlindnera jadinii* and incorporating enzymes on the physical quality of the final aquatic-feed products.

## 2. Materials and Methods Experiment 1

### 2.1 Raw Materials

All raw ingredients presented in Table 1, except *Cyberlindnera jadinii* and fishmeal, were obtained from the Center for Feed Technology, Norwegian University of Life Sciences, Ås, Norway. Inactivated *Cyberlindnera jadinii* yeast was obtained from Lallemand, Salutaguse, Estonia in powder form with a dry matter of 970 g/kg, ash 78 g/kg, crude protein 470 g/kg, crude fat 16 g/kg, and gross energy 19.9 MJ/kg. Fishmeal was obtained from Norsildmel AS, Egersund, Norway with dry matter 917 g/kg, ash 145 g/kg, crude protein 684 g/kg, crude fat 73 g/kg, and gross energy of 19.4 MJ/kg. All materials used for the experimental diets, except *Cyberlindnera jadinii* and vitamin/mineral premix, were milled through a 1 mm screen size installed on an Alpine mill (model 160 UPZ, 1988, No. 13580.1). Yeast and vitamin/mineral premix were not milled due to their particle size below 1mm.

**Table 1.** Formulation of the experimental aquatic feed model presented as the weight (g) and in accordance with the percentage of yeast inclusion.

| *Raw materials ID | *Cyberlindnera jadinii* (CJ) inclusion level (%) |  |  |  |  |
| --- | --- | --- | --- | --- | --- |
|  | 0 CJ | 2.5 CJ | 5 CJ | 10 CJ | 20 CJ |
| Wheat flour | 30 | 30 | 30 | 30 | 30 |
| Fishmeal | 22.5 | 20 | 17.5 | 12,5 | 2.5 |
| Soybean meal | 10 | 10 | 10 | 10 | 10 |
| Poultry meal | 8.5 | 8.5 | 8.5 | 8.5 | 8.5 |
| Rice flour | 6 | 6 | 6 | 6 | 6 |
| Soyprotein concentrate | 6 | 6 | 6 | 6 | 6 |
| Squid meal | 5 | 5 | 5 | 5 | 5 |
| *Cyberlindnera jadinii* | - | 2.5 | 5 | 10 | 20 |
| Monocalcium phosphate | 1.6 | 1.6 | 1.6 | 1.6 | 1.6 |
| Magnesium oxide | 0.3 | 0.3 | 0.3 | 0.3 | 0.3 |
| Mangandioxid | 0.01 | 0.01 | 0.01 | 0.01 | 0.01 |
| Monosodium phosphate | 0.56 | 0.56 | 0.56 | 0.56 | 0.56 |
| Vit/Min premix** | 0.5 | 0.5 | 0.5 | 0.5 | 0.5 |
| Water content (%) | 9 | 9 | 9 | 9 | 9 |
| **TOTAL (%)** | **100** | **100** | **100** | **100** | **100** |

\*Batch size 300 gram; \*\*DSM - OVN™

### 2.2 Preparing the experimental mixes

Mixing of the raw materials was done using a mixer Diosna P1/6 (DIOSNA Dierks & Söhne GmbH, Osnabrück, Germany), with mixing tools based on three agitating paddles and a tulip-form knife. The speed of the paddles during mixing was 250 rpm. and the knife 500 rpm. During the mixing of the formulated diets, 10% of distilled water was added into the mixing mash with a spraying nozzle, model 970, Düsen-Schlick GmbH (Untersiemau/Coburg, Germany). Water was added to integrate enough moisture for further processes. All diets were mixed thoroughly for 800 seconds. From each mixed diet, three samples were randomly taken and thereafter mixed to obtain a representative homogeneous sample. Representative samples were analyzed for moisture content in triplicate by the EU, No. 152/2009 method. The average moisture of the samples for all the trials before the trials was 13 % (+/-1 % w/w). After moisture analyses were performed the mixed mash was vacuum-packed to avoid moisture loss and stored at room temperature of 18°C. Each package was opened right before the pelleting trial.

### 2.3 Steam conditioning and pelleting

To ensure uniformity in size and weight, 0.13 g of feed mash was weighed for each pellet and placed into thirty (30) Eppendorf tubes per trial. Subsequently, all sealed Eppendorf tubes containing the specific feed were immersed in boiling water at atmospheric pressure. The feed mash in the Eppendorf tube was conditioned for 3 minutes in boiling water to allow the water in the feed mash to become steam. Subsequently, the Eppendorf tubes were placed in the fridge at 4 ℃ to cool down for 20 minutes before pelleting so that water could condense swiftly within the entire feed sample in the Eppendorf tube after conditioning. The single-die pelleting method described by Salas-Bringas (2010) was used for the compaction of the mixed feed into cylinder-shaped pellets.

Steam-conditioned material from each Eppendorf tube was used to produce a single pellet by pouring it into a blank die channel preheated to 81°C. The 81°C temperature of the pelleting die is recommended to eliminate potential salmonella contamination following Norwegian law (VKM, 2006). Poured feed mash was heated for 3 minutes before compaction to ensure an equal temperature of the mash particles. Pelleting was done with a compression rod having a diameter of 5.45 mm. The same pelleting temperature was used during material compaction and the extraction of the pellet from the die. During pelleting, an initial pre-load pressure of 240 kPa was used. The setting for a maximal force load for each pellet was 285 N and applied compressibility was 12 MPa. The compacting force load was chosen according to the densities of commercial animal feed pellets produced on ring-die pellet press. This is explained in detail by Salas-Bringas et al. (2011) and Salas-Bringas et al. (2015). The compaction was done at a speed of 10 mm/min with a compression rod inserted in a 5.5-mm die channel with a closed end. Successively after reaching the maximal force load, the closed part of the die was removed. Subsequently, the pellet was discharged from the pelleting die at a pace of 2 mm/min. The selected pace was set to avoid getting beyond the compacting forces and thus avoid additional compaction. The total time of the material retaining in the pelleting channel was 9 minutes. Compacted pellets were 5.5 mm in diameter and had about 0.1 grams of weight. The loss of weight was due to high pressures and temperatures causing moisture evaporation. Pellets were stored at 4 °C, each in the Eppendorf tube prior to further analysis.

## Experiment 2

### 2.4 Preparing the experimental mixes and sampling

The remaining mixed feed from Experiment 1, which was vacuum-packed and stored at room temperature, was used. Each diet was then individually poured into the mixer Diosna P1/6 (DIOSNA Dierks & Söhne GmbH, Osnabrück, Germany). Before mixing, the enzymatic cocktail was prepared with buffer solution and enzymes, non-commercial protease (AB Vista, Marlborough, UK) derived from *Fusarium equiseti,* and endo-exo 1,3-β-glucanase produced by Megazyme, Ireland and derived from *Trichoderma sp.*. Pre-prepared 100 mmol buffer solution (pH 5.5), was stored overnight at 4℃ and used as a carrier for the enzymes. About 0.9 ml of protease and 0.75 ml of endo-exo 1,3-β-glucanase were added to 15 ml of buffer solution. The pH of the mixture was 5.8, close to the producer’s recommendation of pH 5.5 for both enzymatic products. The enzymatic cocktail of 16.65 ml was sprayed on the feed mash batch of 150 g for each feed mixture. The control diet was sprayed only with 16.65 ml of distilled water without enzymes. Spraying was done with a spraying nozzle (Model 970, Düsen-Schlick GmbH, Germany) built into the mixer Diosna P1/6 (Osnabrück, Germany). During the spraying of the enzymatic solution, the mixer was operated with the speed of the paddles set at 250 rpm. The speed of the knife was 500 rpm. All diets were mixed thoroughly for 800 seconds.

Three samples were randomly taken from each mixed diet and subsequently combined to create a representative homogeneous sample. The representative samples were used to analyze moisture content in triplicate for each enzymatically enriched feed mash (EU method No. 152/2009). The average moisture content for all feed diets was 15% (0.3% w/w). The remaining treated feed mash was removed from the mixer, vacuum-packed, and stored at 4°C to preserve moisture content until further steam conditioning and pelleting. Steam conditioning and pelleting were done in the same way as in experiment 1.

## 3. Analytical techniques and measurements

### 3.1 Particle size distribution

The laser diffraction method was used to verify the particle size distribution of the feed-mixed mash. The device used for measuring particle size distribution was Mastersizer S instrument (Malvern Instruments Ltd, Worcestershire, UK). Calculating the volumetric particle size distribution of the light energy on the detector was done by deflecting the light based on the theory for spherical particles (Allen, 1997).

### 3.2 Measurement of compaction, pellet discharge, and pressure at initial flow (p_max_)

Maximum compaction pressure (Pa) at maximum force (N) during densification, compaction, and pellet ejection from the die were observed and recorded. The recording was done with NEXIGEN Plus software connected to a Lloyd texture analyzer (LR 5K Plus; Lloyd Instruments, U.K.). Measuring maximum compaction pressure at maximum force might point to possible alterations in electrical energy consumption connected to alterations in the resistance of the feed material to move through the die (p_max_). The p_max_ was identified as the pressure needed to initiate pellet ejection from the pelleting die when the close end from the die channel was removed. Measurements were done as a flow resistance indicator to measure variations between trials, caused by friction on the contact area between the die and compacted pellet. The initial movement of the pellets through the die was recorded as a force needed for the pellet to start moving through the open 5.5 mm die hole. Measurements were done after compaction and when the blank part of the die was removed. The pellet discharge speed was set to 2 mm/min to guarantee that the pressure would be under the previous compaction pressures. The analytical results for maximal pressure before the pellet started moving through the die, were obtained using a Lloyd LR 5K Plus texture analyzer and assessed with Eq. 1.

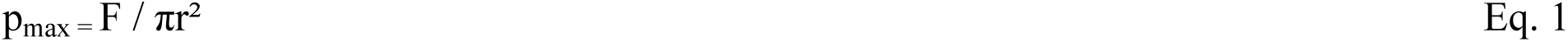

Where:

F – Load force (Nm)

r - radius of a pellet (mm)

### 3.3 Tensile strength

Results of the tensile strength analyses were done by applying maximum force (F) on the compacted feed pellet under diametral compression. Three randomly chosen pellets for each treatment were used. Tensile strength was measured for each pellet by recording the first peak force (F) while compressing the pellet across the diameter at a speed of 1 mm/minˉ. The stress (σ) was estimated by using Eq. 2, the Brazilian test. The Brazilian test is a geotechnical laboratory method used to indirectly measure the tensile strength of materials, by applying diametrical compression to a disc-shaped specimen until it splits. A probe with a surface of 60 mm in diameter was used to measure σ (Eq. 2) while being connected to a Lloyd LR 5K Plus texture analyzer (Lloyd Instruments, U.K.)

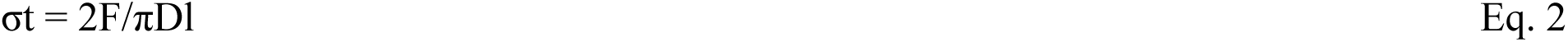

Where:

σt - maximum tensile strength (MPa)

F - load at fracture of the first peak force (N) D – shrimp feed pellet diameter (mm)

l - shrimp feed pellet length (mm)

### 3.4 Measuring contact surface angle (θ)

The contact surface angle measures how a liquid droplet interacts with a solid surface, indicating wettability. Droplets are used because their shape reveals the balance between adhesive and cohesive forces. The droplets of oil and water were disposed from a dosing needle and placed at the diametral plain surface of a feed pellet, defined as the upper plain being at the side of a compression rod during pelleting. The analyses of liquid droplets from a dosing needle, defined as contact surface angle, outline the surface energy between the adhering liquid and the pellet surface. Surface wetting of the shrimp pellets, the effect of added yeast, and the enzyme treatment on θ for both oil and water were evaluated with a video-based device OCA 15EC (Data Physics Instruments GmbH, Germany) (Fig. 1). The θ analyses were completed on three randomly chosen pellets to compare how liquid sorption of the pellets with rape seed oil or distilled water could be affected by the experimental factors. The droplet volume for distilled water was 1 μl and for rapeseed oil 5 μl. Due to the larger surface tension of rapeseed oil, it was more difficult to detach the oil droplet from the needle, thus a larger droplet volume was needed. The recorded droplet absorption is measured in seconds and presented in the results as the initial surface angle (T0) and final droplet age. Rapeseed oil θ and distilled water θ were recorded within 3 and 1.2 seconds, respectively. The time frame was decided for both oil and water droplets by full penetration of the liquids at the upper diametral plain surface of a shrimp pellet. The θ measurements were done at a temperature of 18 °C. The software SCA20 (Dataphysics instruments GmbH, Germany) was used to record the change of θ at different time intervals. When the initial θ was < 90°, the surface was considered as hydrophilic, however, if θ > 90° a hydrophobic surface was considered, described by Förch et al. (2009) and Mišljenović et al. (2015).

**Figure 1.**
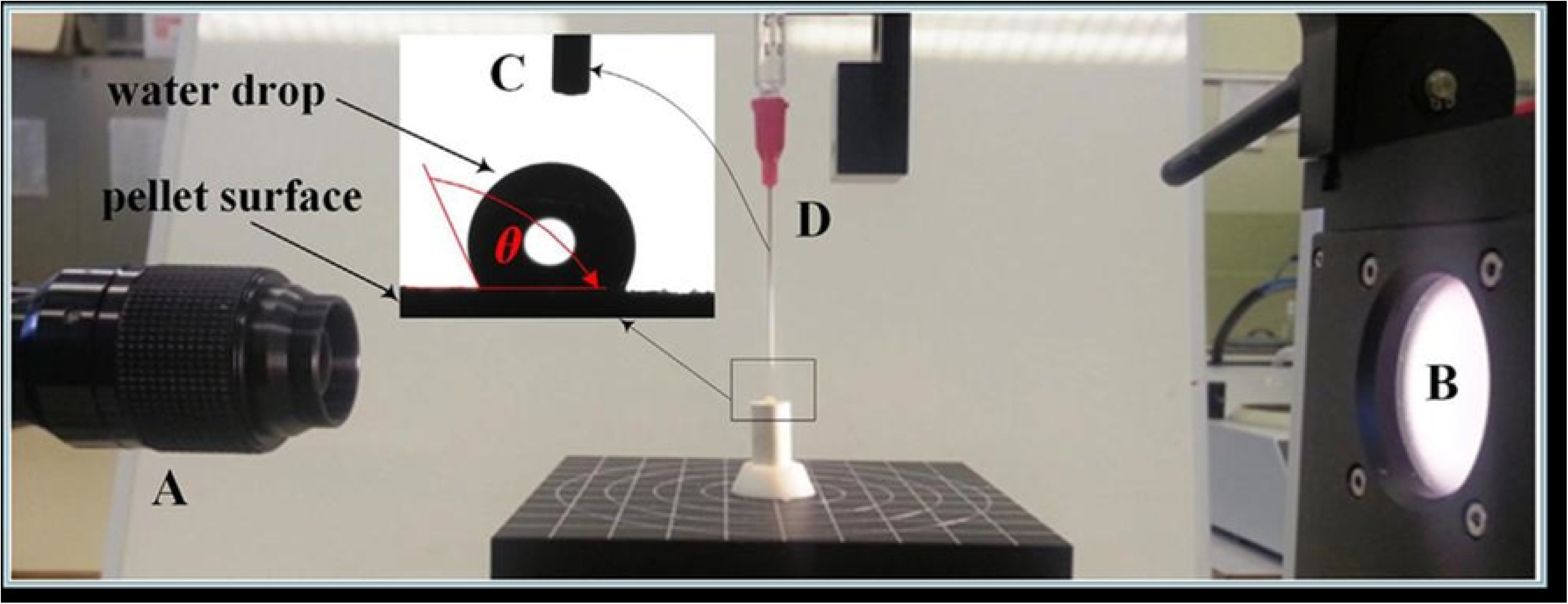
Setup for surface angle measurement. Letters indicating: A - video camera; B – light source; C – an image of a drop on top of a pellet surface; D – dosing syringe. Adopted from Mišljenović et al. (2015).

### 3.5 Underwater pellet swelling rate analyses (UPS)

The UPS measurements explain if increasing yeast content in the feed instead of fishmeal and enzymatic treatments could influence changes in the physical properties of the pellets. Also, the UPS indicates pellet cohesivity (particle detachment), linked to the swelling rate of the pellet. A special prearrangement to monitor the swelling of shrimp feed pellets underwater was used as described by Miladinovic et al. (2021). The constructed system consisted of a video microscope (Microviper) and microscope lens (Allen ¼”) mounted on the optical tensiometer (OCA 15EC, Data Physics Instruments GmbH, Germany) as shown in Fig. 2 was used for the UPS measurements. Optical monitoring of pellet swelling was done in accordance with Ferreira and Rasband (2012) with the image processing software Fiji. Distilled water at a temperature of 20 °C was added to the experimental glass container. For each treatment, four randomly chosen pellets were used for measuring UPS. The reference of a starting point for UPS measurement was a cross-sectional view of the feed pellet with a diameter of 5.5 mm. An image of a cross-sectional view of the feed pellet was taken every 60 seconds for an observation time of 40 minutes. The length of the observation time was chosen as the time required for shrimp to consume the pellet in the open-pond shrimp farming system (Lovell, 1998). The observation area defining the cross-section of a swollen shrimp feed pellet underwater during observation time implies the UPS results.

**Figure 2.**
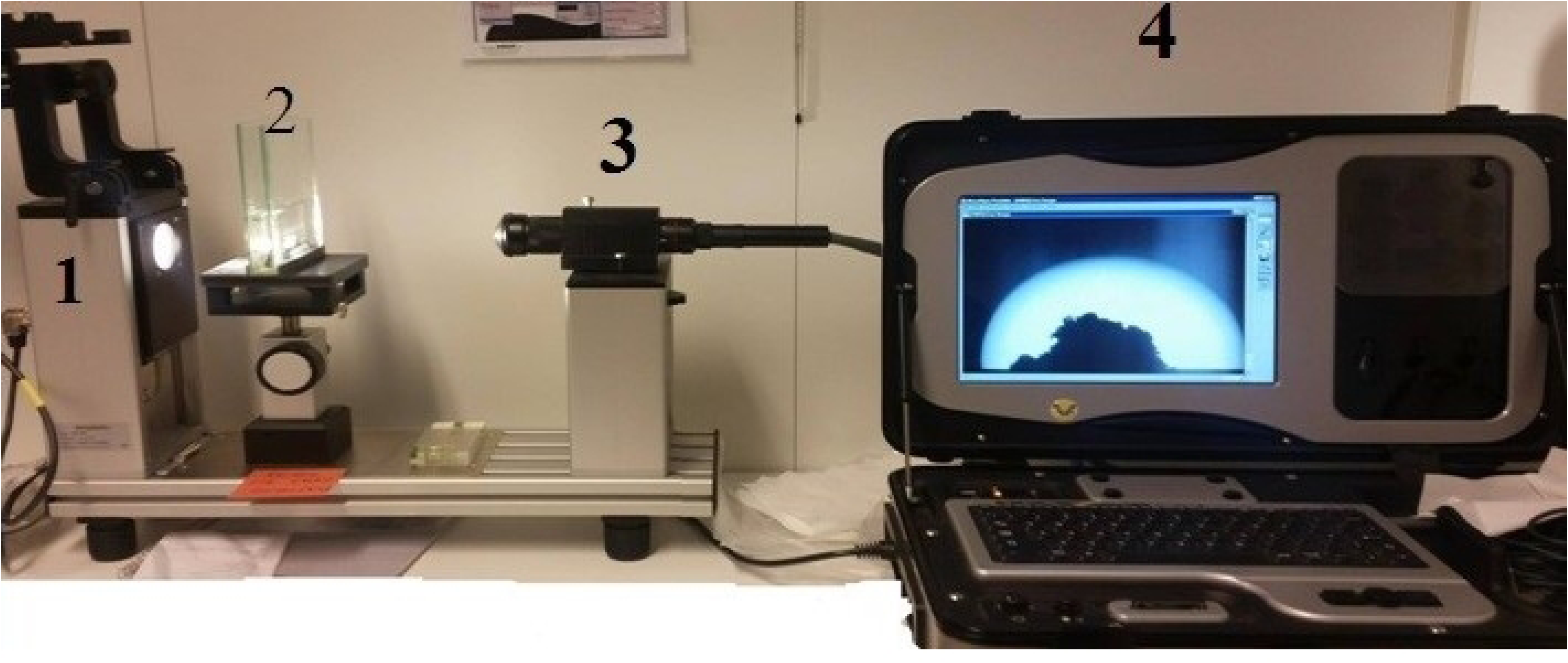
Full setup instrument for the UPS measurements (1 - Krüss Tensiometer; 2 - Water glass container with the pellet; 3 - Allen zoom compact video microscope lenses; 4 - Microviper portable computer)

### 3.6 Surface roughness analyses

Surface roughness analyses were done to understand if the replacement of the fish meal with yeast or the addition of enzymes may influence any changes to particle packing and thus surface characteristics of the pellets. Results are represented by the irregularities existing on the surface of the feed pellets in the diametral and longitudinal directions. The diametral surface roughness has been created by interactions between feed particles during compaction. However, longitudinal surface roughness is created as the interaction between particles and the die wall. Surface roughness is further presented as a numerical scale of the surface condition influenced by novel materials, enzymes, and diverse packing-ability of the particle in the pellets. The analyses were done with a surface roughness tester (Surftest SJ-210, Mitutoyo, Japan).

### 3.7 Data analyses

The software used for statistical analyses was Minitab v.17 and for plotting the figures was utilized Microsoft Excel. ANOVA analyses were used to examine the possible effects of the increased percentage of yeast and enzyme addition. Tukey–Kramer method, using a 95 % confidence interval, was used to show significant differences between treatments. Pearson correlation test with a 95 % confidence interval was used to analyze correlations between variables.

## 4. Results

**Experiment 1 – replacement of the fish meal with *Cyberlindnera jadinii***

### 4.1 Particle size of the mash prior to compaction

All diets with yeast inclusion had similar particle size distribution, where large particles were between 400 μm and 500 μm. Medium size particles were found to be between 80 and 120 μm with a significant decrease when adding more yeast to the feed. The same was observed with the smallest particles under 20 μm (Fig. 3).

**Figure 3.**
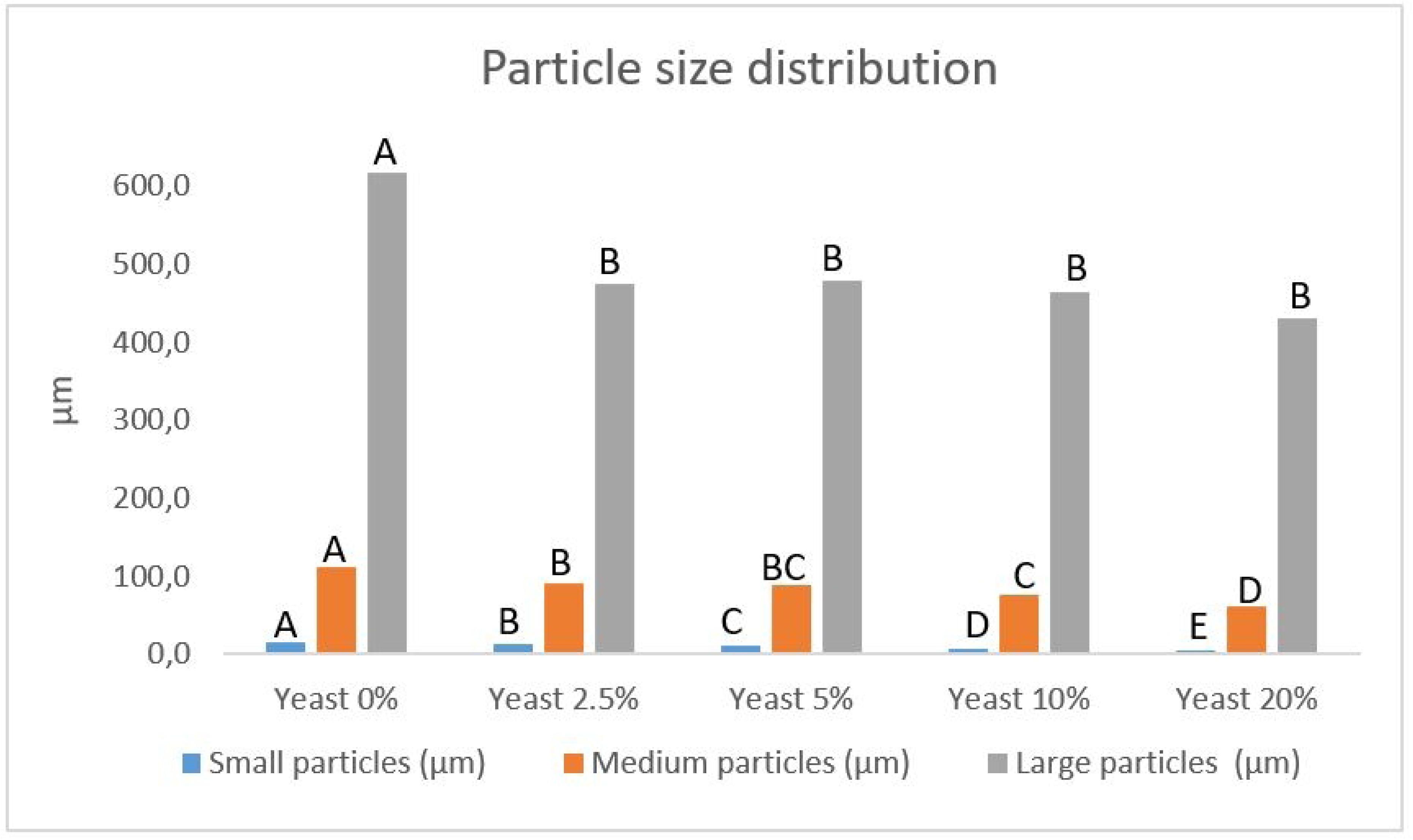
Particle size distribution of the shrimp feed diets. Data representing means based on two replications. Different sequential letters (A, B, C, D, E) from the ANOVA-Tukey statistical method indicate differences at a 5% level

### 4.2 Flow resistance in the die during discharge of the shrimp feed pellet (p_max_)

The p_max_ results presented in Table 2 do not show any distinctive change when yeast is added and compared to control feed without added yeast. However, pelleting only yeast as a negative control, increased p_max_ over 17 folds compared to all other diets (p<0.05). Adding from 2.5% to 20% yeast did not change p_max_.

**Table 2.** Experiment 1. Results of p_max_; a_w_ - water activity; tensile strength; oil and water contact angle (CA) and underwater pellet swelling (UPS) Data representing means and ±SD. Different sequential letters from ANOVA-Tukey statistical method signify the difference at 5% level. Presented p-values for p-max (p<0.001); a_w_ (p>0.05); Tensile strength (p<0.001); CA-oil (p<0.05); CA-water (p<0.01) and UPS (p<0.05)

| Added yeast | p-max (MPa) | aW | Tensile strength (N/mm<sup>2</sup>) | CA - oil at 0 sec. | CA - oil at 47 sec. | CA - oil at 94 sec. | CA - water at 0 sec. | CA - water at 47 sec. | CA - water at 94 sec. | UPS (mm<sup>2</sup>) at 1 min. | UPS (mm<sup>2</sup>) at 20 min. |
| --- | --- | --- | --- | --- | --- | --- | --- | --- | --- | --- | --- |
| 0 % | 18.2<sup>b</sup> ±6.0 | 0.35<sup>a</sup> | 18.1<sup>c</sup> ±1.5 | 64.0<sup>a</sup> ±2.0 | 62.9<sup>a</sup> ±2.1 | 61.6<sup>a</sup> ±2.2 | 63.1<sup>a</sup> ±1.3 | 61.3<sup>a</sup> ±1.5 | 59.5<sup>a</sup> ±2.2 | 30.3<sup>ab</sup> ±0.01 | 40.2<sup>ab</sup> ±7.5 |
| 2.5% | 16.6<sup>b</sup> ±1.0 | 0.46<sup>a</sup> | 34.1<sup>c</sup> ±14.8 | 63.3<sup>ab</sup> ±1.5 | 60.7<sup>ab</sup> ±1.3 | 58.8<sup>ab</sup> ±1.0 | 59.7<sup>ab</sup> ±4.4 | 53.2<sup>ab</sup> ±5.1 | 47.9<sup>b</sup> ±5.7 | 36.1<sup>a</sup> ±1.8 | 40.8<sup>ab</sup> ±3.5 |
| 5 % | 17.4<sup>b</sup> ±3.6 | 0.35<sup>a</sup> | 30.7<sup>c</sup> ±3.1 | 57.5<sup>ab</sup> ±10.1 | 54.2<sup>ab</sup> ±10.7 | 57.4<sup>ab</sup> ±4.9 | 59.1<sup>ab</sup> ±5.1 | 49.3<sup>ab</sup> ±6.5 | 43.7<sup>b</sup> ±5.5 | 28.4<sup>ab</sup> ±6.9 | 49.1<sup>ab</sup> ±5.7 |
| 10 % | 19.7<sup>b</sup> ±4.1 | 0.40<sup>a</sup> | 44.2<sup>bc</sup> ±6.6 | 61.2<sup>ab</sup> ±3.0 | 59.3<sup>a</sup> ±3.0 | 57.6<sup>bc</sup> ±3.5 | 51.0<sup>b</sup> ±3.8 | 44.0<sup>bc</sup> ±4.5 | 40.1<sup>bc</sup> ±3.4 | 33.0<sup>ab</sup> ±2.1 | 47.2<sup>ab</sup> ±10.8 |
| 20 % | 23.9<sup>b</sup> ±5.7 | 0.40<sup>a</sup> | 70.3<sup>b</sup> ±5.0 | 50.7<sup>ab</sup> ±1.2 | 49.6<sup>bc</sup> ±2.1 | 48.7<sup>bc</sup> ±2.7 | 49.9<sup>bc</sup> ±5.7 | 44.9<sup>bc</sup> ±4.7 | 42.5<sup>bc</sup> ±4.6 | 28.8<sup>ab</sup> ±0.8 | 51.0<sup>a</sup> ±2.1 |
| 100 % | 128.7<sup>a</sup> ±43.1 | 0.35<sup>a</sup> | 173.8<sup>a</sup> ±29.7 | 49.4<sup>b</sup> ±7.0 | 48.1<sup>c</sup> ±6.7 | 47.00<sup>c</sup> ±6.4 | 38.3<sup>c</sup> ±4.3 | 33.2<sup>c</sup> ±3.5 | 31.0<sup>c</sup> ±2.7 | 25.1<sup>b</sup> ±0.5 | 32.8<sup>b</sup> ±0.7 |

Data representing means and ±SD. Different alphabetical letters in superscripts from ANOVA-Tukey statistical method indicate significant difference at 5% level. Presented p-values for p-max (p<0.001); aW (p>0.05); Tensile strength (p<0.001); CA-oil (p<0.05); CA-water (p<0.01) and UPS (p<0.05)

### 4.2 Tensile strength of pellets

The hardness of pellets measured as tensile stress increased significantly (p<0.05) by including 20% yeast in the feed when compared to the control diet and the diets with 2.5% and 5% yeast. Pelleting only yeast without other ingredients showed increased hardness of the pellets by over 9 folds (Table 2). Tensile strength showed to be moderately correlated to p_max_ (Fig. 4), where almost half of the tensile strength as a dependent variable may be explained by flow resistance during discharge of the shrimp feed pellet from the die (R^2^ = 0.49).

**Figure 4.**
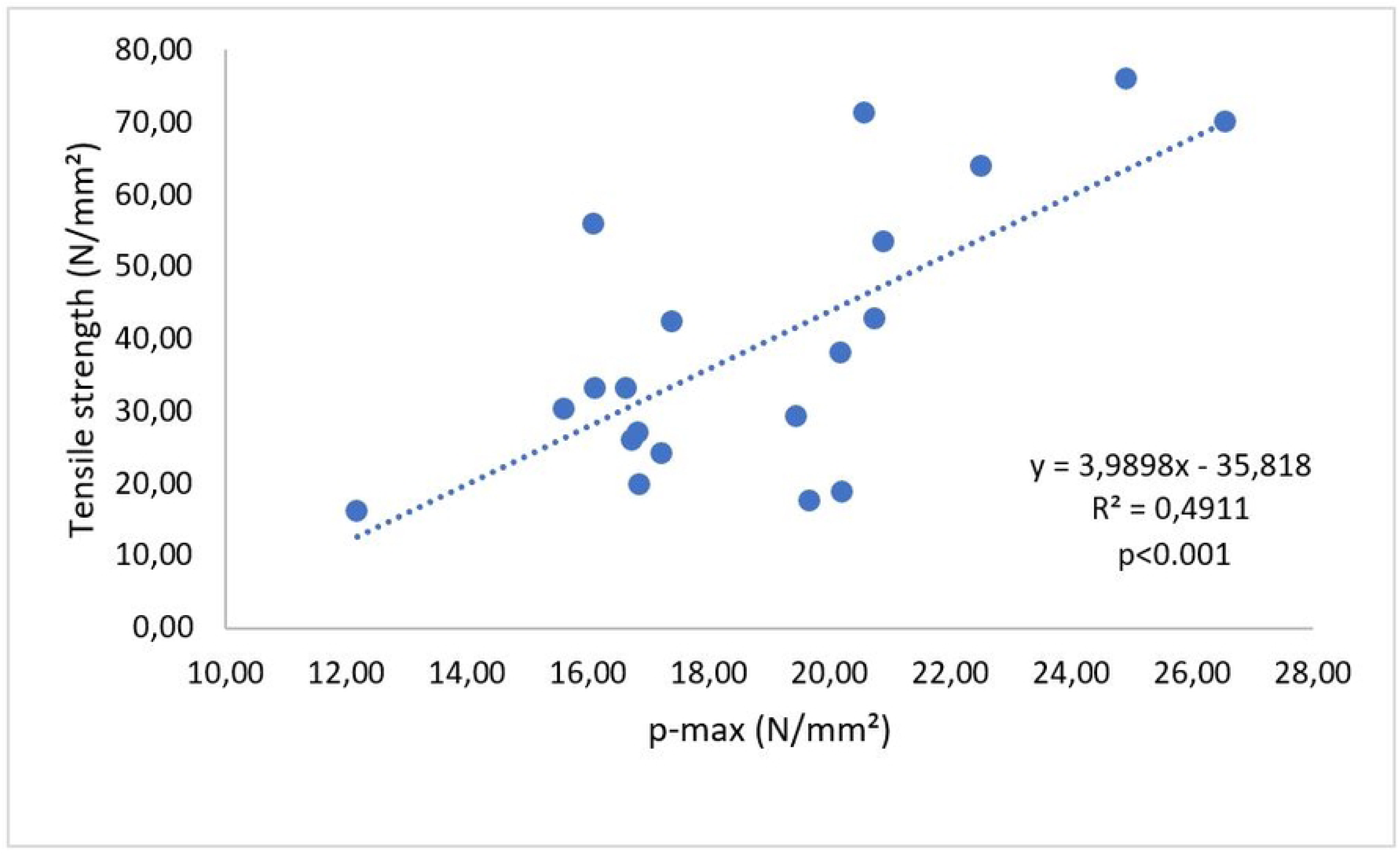
Correlation between p_max_ and tensile strength in experiment 1 (p<0.001)

### 4.3 Surface contact angle (θ) measurements for oil and water

By adding 2.5% and up to 20% yeast in the feed, the θ of the oil at zero time was not different from that of the control feed. A difference at zero time for oil θ was, however, observed between 100% yeast and 0% yeast (p<0.05). Pellets made of 100% yeast were more lipophobic as compared to pellets with no added yeast. At 47 seconds the lipophilicity was more pronounced (p<0.05) for feed pellets without yeast as compared to feed with 20% yeast and 100% yeast. Similar results for oil θ were observed when pellets with no added yeast were compared with 10%, 20%, and 100% yeast at 94 seconds. Pellets with 10%, 20%, and 100% yeast at time zero showed to have pronounced aquaphobic behaviour when compared to control pellets without added yeast. Similar effects were observed at 47 seconds. However, at 94 seconds the water θ showed to be significantly lower (p<0.01) in the pellets with from 2.5% and up to 100% yeast (Table 2).

### 4.4 Underwater pellet swelling (UPS)

The UPS did not differ when control pellets (0% yeast) were compared to other pellets at the first minute after submersion under water. Differences were observed between pellets with 2.5% and 100% yeast at minute 1. Pellets containing 100% yeast tended to swell poorly at the first minute. After 20 minutes the pellets with 20% yeast had swollen 1.8 folds and the pellets with 100% yeast about 1.3 folds. Pellets with 100% yeast showed significantly (p<0.05) slower swelling when compared to pellets with 20% added yeast (Table 2). UPS results showed having low correlation to tensile strength (R² = 0.3) in experiment 1.

### 4.6 Surface roughness at diametral and longitudinal direction

Irregularities at the surface of the pellets defined as surface roughness are presented in Fig. 5. Diametral surface roughness was different when yeast was added to the feed as compared to the control feed with no yeast. However, no significant change in surface roughness was observed for the diet with 5% yeast. Longitudinal roughness difference was observed only for the pellets with 20% yeast in comparison with the control pellets, and pellets with 2.5% or 100% yeast. A correlation between p_max_ and longitudinal surface roughness was observed. More than half of the longitudinal surface roughness, as a dependent variable, may be explained by flow resistance in the pelleting die during shrimp feed pellet discharge (R^2^ = 0.51) (Fig. 6). However, the correlation between the diametral surface roughness and p_max_ was not observed (R^2^ = 0.07).

**Figure 5.**
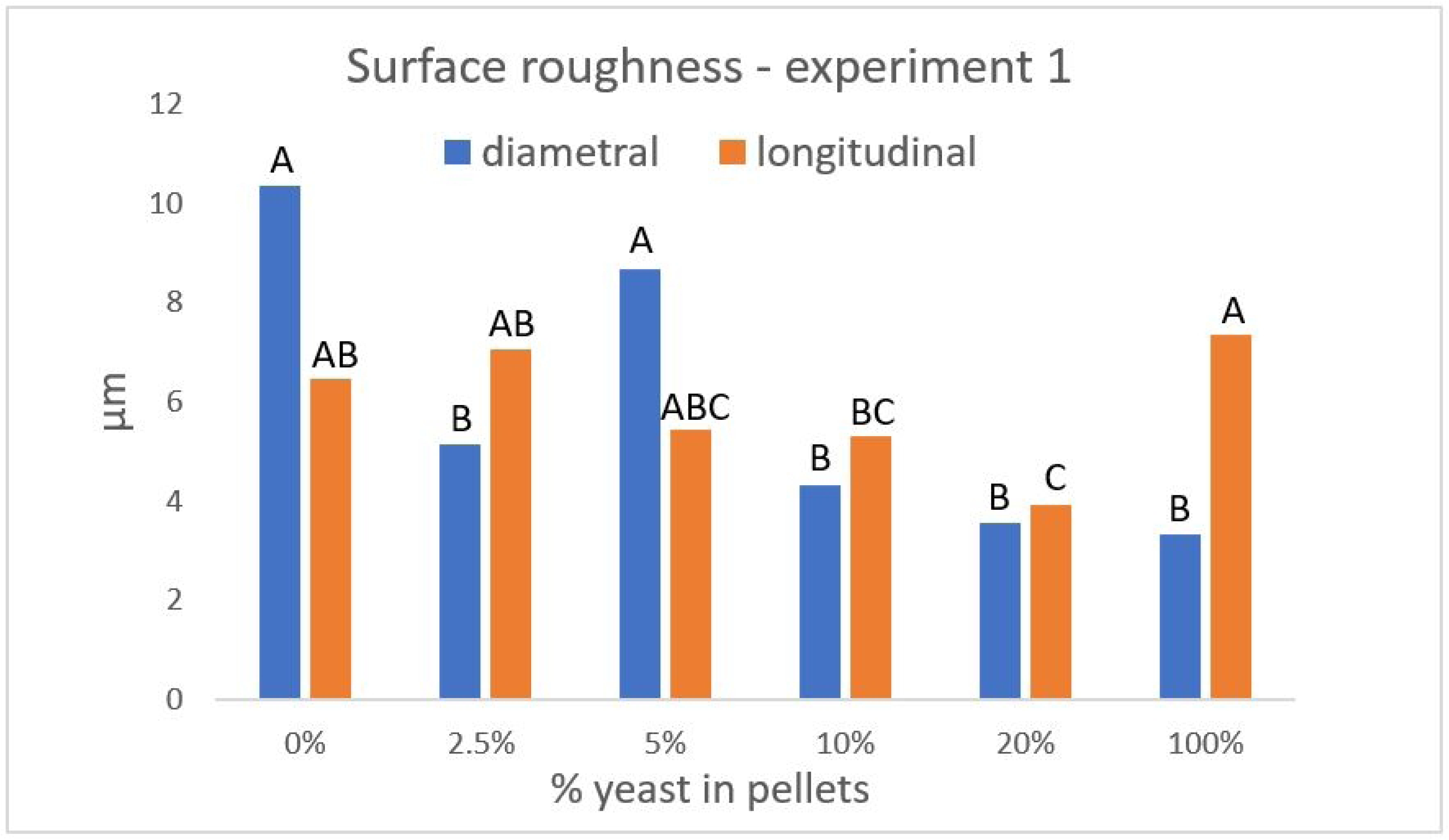
Surface roughness of the pellets with no added enzymes for both diametral and longitudinal analyses. Data representing means based on 25 repetitions. Different sequential letters (A, B, C) from ANOVA-Tukey statistical method signify difference at a 5% level separately for diametral and for longitudinal measurements where p-values were lower than 0.001.

**Figure 6.**
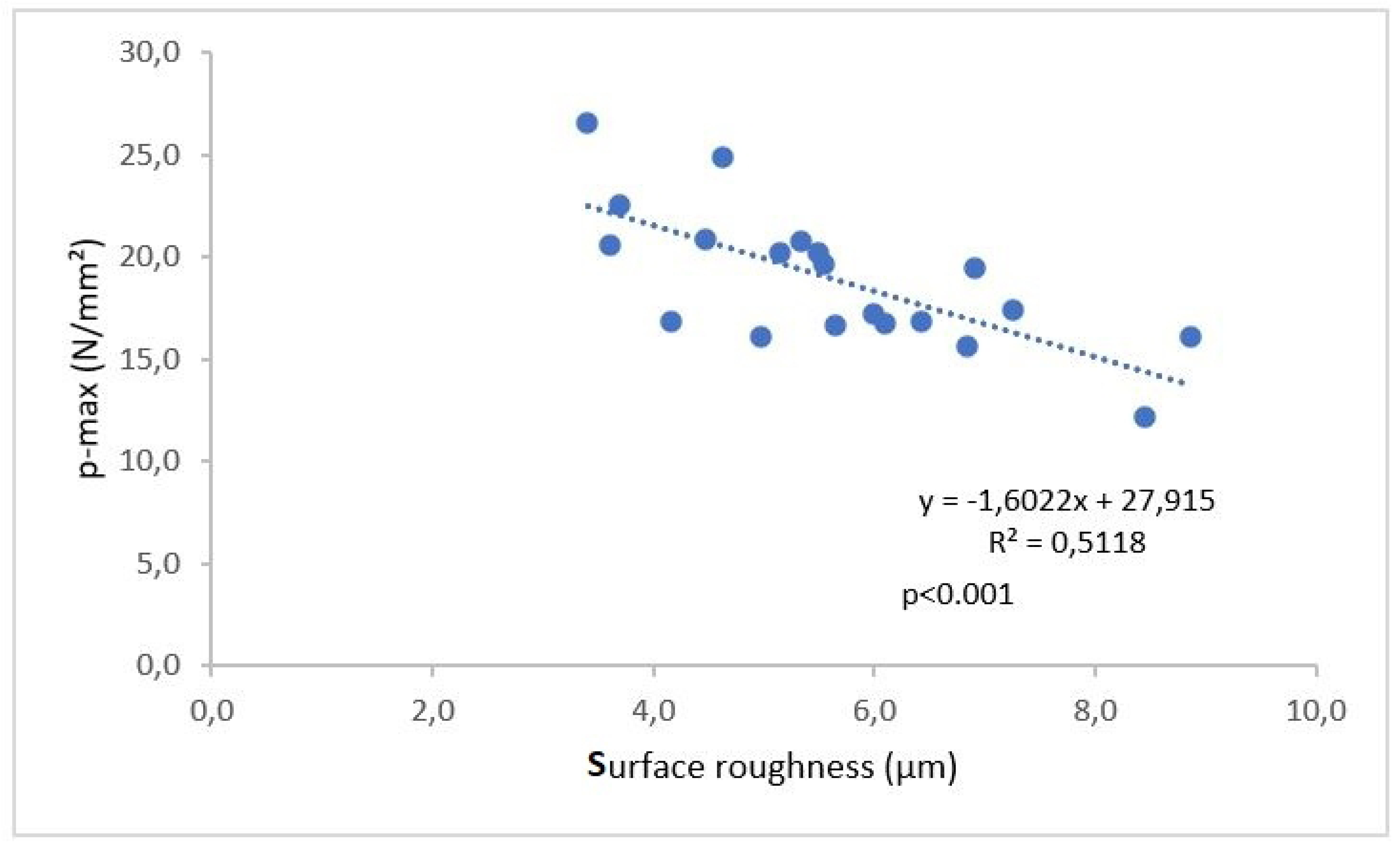
Correlation between longitudinal surface roughness and p_max_ based on % of added yeast from 0 to 100 %

**Experiment 2 – Enzymatic treatment of the feed and its effect on physical pellet properties**

### 4.7 Flow resistance in the die during discharge of the shrimp feed pellet (p_max_)

By adding enzymes in the feed with the inclusion of up to 5% yeast, no change in p_max_ was detected. However, when enzymes were included in the feed with 10% and 20% yeast a significant increase in p_max_ was identified (p<0.001) when compared to the control feed with added enzymes (Table 3).

**Table 3.** Experiment 2. Results of p_max_; a_w_ - water activity; tensile strength; oil, and water contact angle (CA); and underwater pellet swelling (UPS) Data representing means and ±SD. Different letters from ANOVA-Tukey statistical method indicate significant differences at 5% level. Presented p-values for p-max (p<0.01); a_w_ (p<0.05); Tensile strength (p<0.001); CA-oil (p<0.01); CA-water (p<0.01) and UPS (p<0.01)

| Experiment 2 |  |  |  |  |  |  |  |  |  |  |  |
| --- | --- | --- | --- | --- | --- | --- | --- | --- | --- | --- | --- |
| Added yeast | p-max (MPa/mm<sup>2</sup>) | aW | Tensile strength (N/mm<sup>2</sup>) | CA - oil at 0 sec. | CA - oil at 5 sec. | CA - oil at 10 sec. | CA - oil at 15 sec. | CA - oil at 20 sec. | CA - water at 0 sec. | CA - water at 0.5 sec. | CA - water at 1 sec. |
| 0% no enz | 0.4<sup>bc</sup> ±0.1 | 0.46<sup>ab</sup> ±0.01 | 9.3<sup>bc</sup> ±3.1 | 53.4<sup>a</sup> ±0.5 | 42.2<sup>a</sup> ±0.6 | 41.3<sup>a</sup> ±0.6 | 38.5<sup>a</sup> ±0.3 | 38.0<sup>a</sup> ±0.3 | 53.1<sup>a</sup> ±2.5 | 46.5<sup>a</sup> ±3.0 | 40.4<sup>a</sup> ±2.3 |
| 0% + enz. | 0.3<sup>c</sup> ±0.08 | 0.46<sup>ab</sup> ±0.01 | 6.3<sup>c</sup> ±1.8 | 52.1<sup>ab</sup> ±0.2 | 41.5<sup>a</sup> ±0.6 | 37.7<sup>b</sup> ±0.6 | 35.2<sup>b</sup> ±0.3 | 31.7<sup>b</sup> ±0.3 | 58.1<sup>a</sup> ±1.1 | 45.4<sup>ab</sup> ±1.2 | 37.9<sup>a</sup> ±1.6 |
| 2.5% + enz. | 0.3<sup>bc</sup> ±0.04 | 0.45<sup>ab</sup> ±0.01 | 8.3<sup>bc</sup> ±0.7 | 51.0<sup>b</sup> ±0.5 | 41.4<sup>a</sup> ±0.4 | 36.7<sup>c</sup> ±0.6 | 33.9<sup>b</sup> ±0.2 | 31.6<sup>bc</sup> ±0.3 | 59.0<sup>a</sup> ±3.1 | 39.7<sup>ab</sup> ±1.9 | 32.2<sup>ab</sup> ±3.1 |
| 5% + enz. | 0.3<sup>bc</sup> ±0.02 | 0.44<sup>b</sup> ±0.01 | 7.9<sup>c</sup> ±0.6 | 52.1<sup>ab</sup> ±0.3 | 35.5<sup>b</sup> ±0.6 | 33.5<sup>c</sup> ±0.6 | 30.7<sup>c</sup> ±0.5 | 28.7<sup>d</sup> ±0.4 | 56.2<sup>a</sup> ±1.2 | 29.3<sup>b</sup> ±2.2 | 19.2<sup>b</sup> ±3.6 |
| 10% + enz. | 0.5<sup>b</sup> ±0.01 | 0.48<sup>a</sup> ±0.01 | 12.4<sup>b</sup> ±0.3 | 45.9<sup>c</sup> ±0.1 | 36.1<sup>b</sup> ±0.6 | 32.7<sup>c</sup> ±0.8 | 31.1<sup>c</sup> ±0.2 | 29.7<sup>d</sup> ±0.4 | 57.1<sup>a</sup> ±0.9 | 35.3<sup>ab</sup> ±1.7 | 30.6<sup>ab</sup> ±2.1 |
| 20% + enz. | 0.8<sup>a</sup> ±0.03 | 0.48<sup>a</sup> ±0.01 | 19.2<sup>a</sup> ±0.7 | 46.4<sup>c</sup> ±0.3 | 38.1<sup>b</sup> ±0.6 | 35.1<sup>bc</sup> ±0.6 | 32.0<sup>c</sup> ±0.4 | 29.9<sup>cd</sup> ±0.5 | 58.9<sup>a</sup> ±4.4 | 49.7<sup>a</sup> ±7.0 | 49.7<sup>a</sup> ±7.0 |

### 4.8 Tensile strength of pellets

The tensile strength of the pellets was significantly increased in the feed containing 20% yeast and added enzymes (p<0.001). This increase was not observed in other yeast-containing feeds with added enzymes when compared to the control feed (Table 3). Additionally, it was found that p_max_ had a very strong influence on the tensile strength of the pellets (Fig. 7).

**Figure 7.**
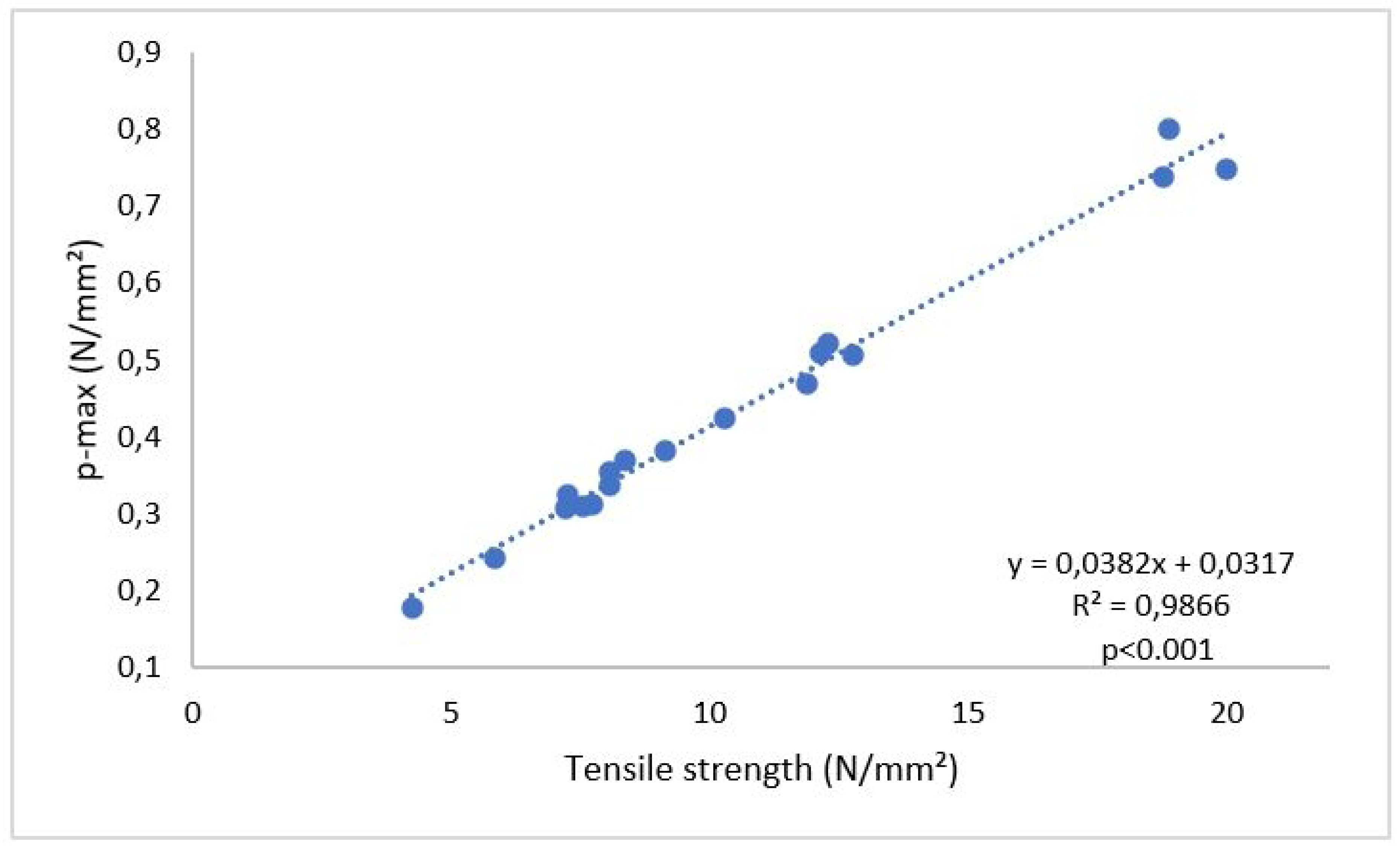
Correlation between p_max_ and tensile strength in experiment 2 (p<0.001) based on % of added yeast whether treated or not treated with enzymes

### 4.9 Surface contact angle (θ) measurements for oil and water (CA)

At zero time, pellets with 10% and 20% yeast and added enzymes showed a lipophobic behaviour (p<0.01) when compared to other treatments (Table 3, Fig. 8). After 5 seconds the lipophobic behaviour was also observed on the surface of pellets containing 5% yeast and added enzymes. At 10 seconds, this was also seen for enzymatically treated pellets containing 2.5% yeast and 0% yeast. A similar response was observed for the rest of the analytical time, i.e. after 15 and 20 seconds (Table 3).

**Figure 8.**
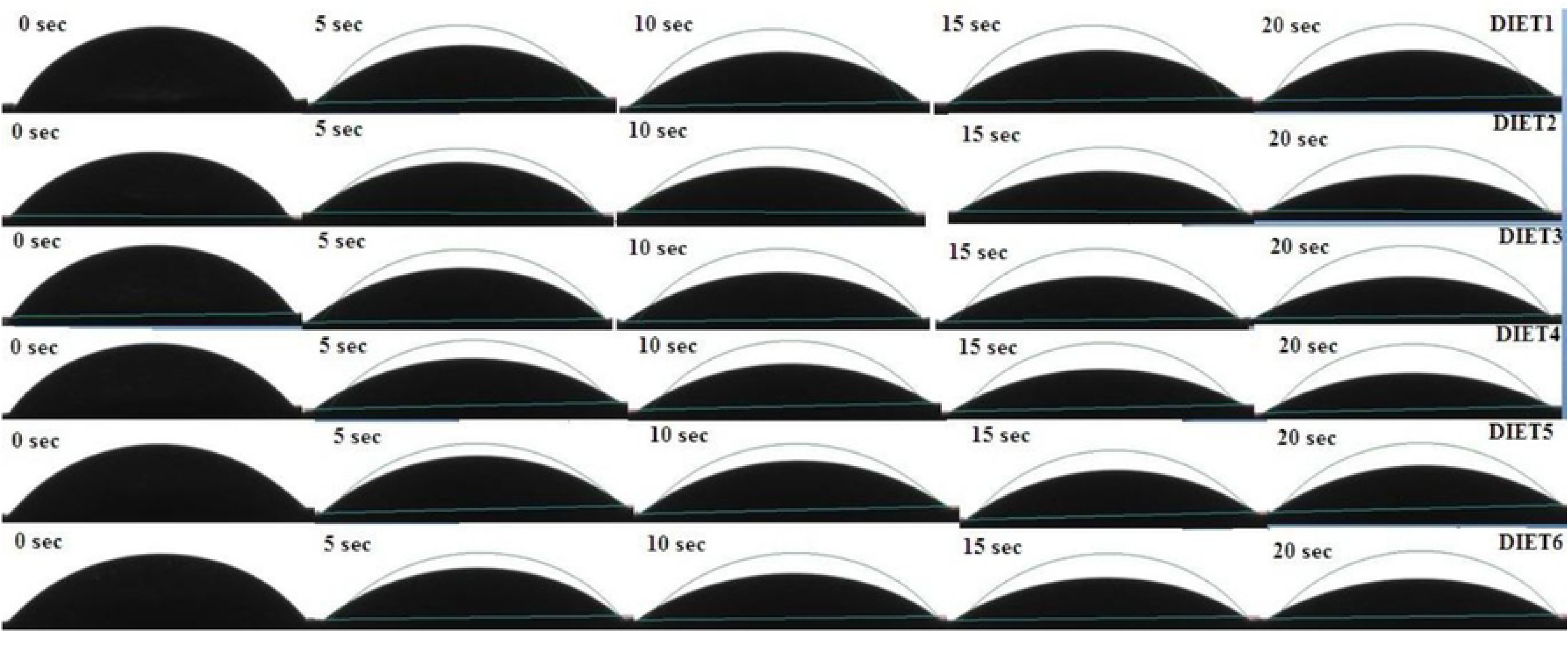
Change of contact angle (CA) from oil drop on the pellet surface at different time intervals. The curved line symbolizes the initial oil drop profile.

CA of the water drop placed at the pellet surface measured every 0.5 seconds for a total measuring time of 2 seconds, showed similar results for all feed at time zero. However, after 0.5 seconds the enzymatically treated pellet with 5% yeast had a significantly lower contact angle when compared to control feed without enzymes and pellets treated with enzymes containing 20% yeast (Table 2, Fig. 9). Similar observations were seen after 1, 1.5 and 2 seconds of CA measurements.

**Figure 9.**
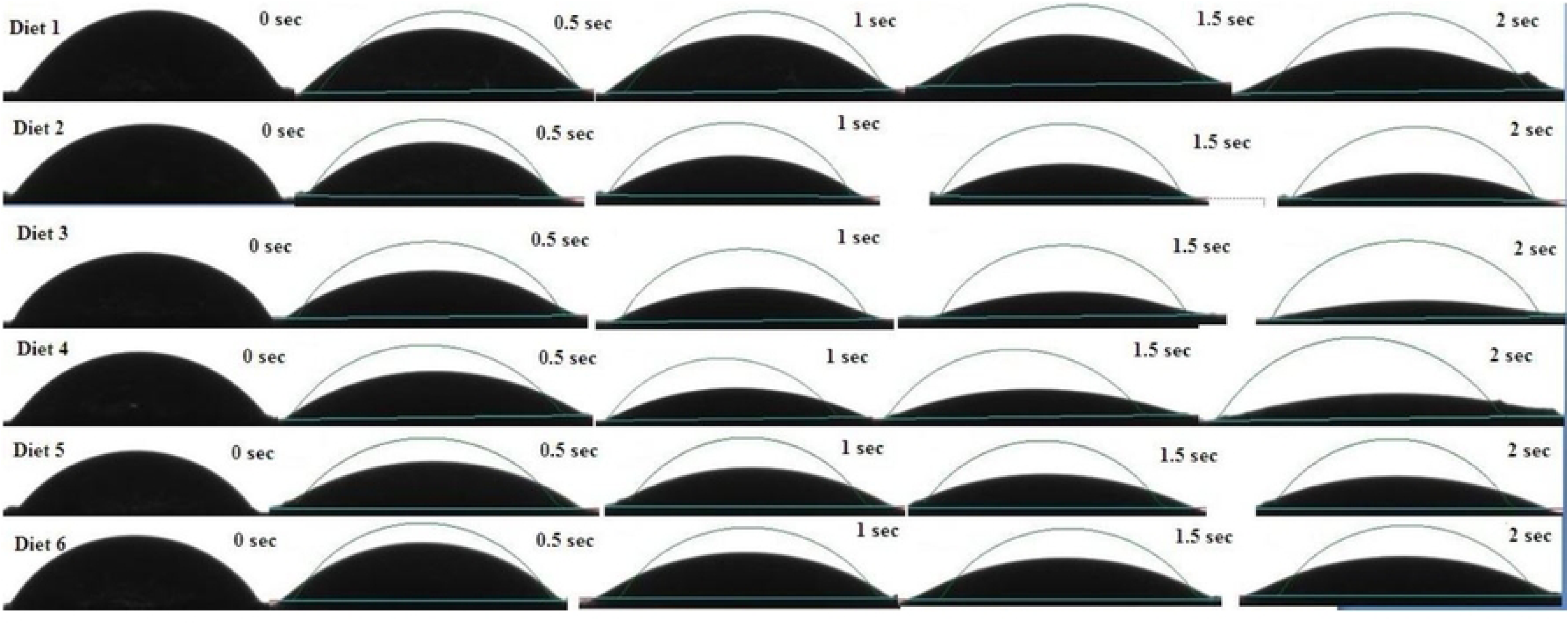
Change of CA from water drop on the pellet surface at different time intervals. The curved line symbolizes the initial water droplet profile.

### 4.10 Underwater pellet swelling rate (UPS)

The UPS analyses showed that by adding enzymes to the control feed without yeast there was a significant increase in pellet swelling one minute after submersion in water. After 40 minutes, the control feed with and without added enzymes was equally swollen. Conversely, the inclusion of 10% and 20% of yeast significantly decreased the UPS when compared to the enzymatically treated control feed (p=0.003) (Table 3). A similar trend was observed after 20 and 40 minutes after submersion in stagnant water. The UPS linearly decreased by increasing yeast addition in the shrimp feed, indicating that the addition of 20% yeast may result in a UPS decrease of about 31%. The UPS of the feed pellets showed to be independent of the tensile strength (R² = 0.24).

### 4.11 Water activity (a_w_)

Differences in a_w_ values were not observed in experiment 2 (Table 3). Thus, the a_w_ analyses showed that the substitution of fishmeal with yeast and added enzymes in shrimp feed did not influence a_w_.

### 4.12 Surface roughness at the diametral and longitudinal direction of pellets treated with enzymes

In this work, irregularities at the surface of pellets compressed with the same forces are defined in this work as surface roughness. Fig. 10 shows that longitudinal surface roughness presented as the interaction between particles and the pelleting die wall is lowered in feed containing 5% and up to 20% yeast and added enzymes. The longitudinal surface roughness results indicate a linear decrease of surface roughness along the longitudinal pellet wall by replacing fishmeal with yeast when the feed mash was treated with enzymes. However, for diametral surface roughness, created by interactions between feed particles during compaction, the same can be concluded only for 5% and 10% added yeast. No significant difference in diametral surface roughness between enzyme-treated feed containing 2.5% or 20% yeast.

**Figure 10.**
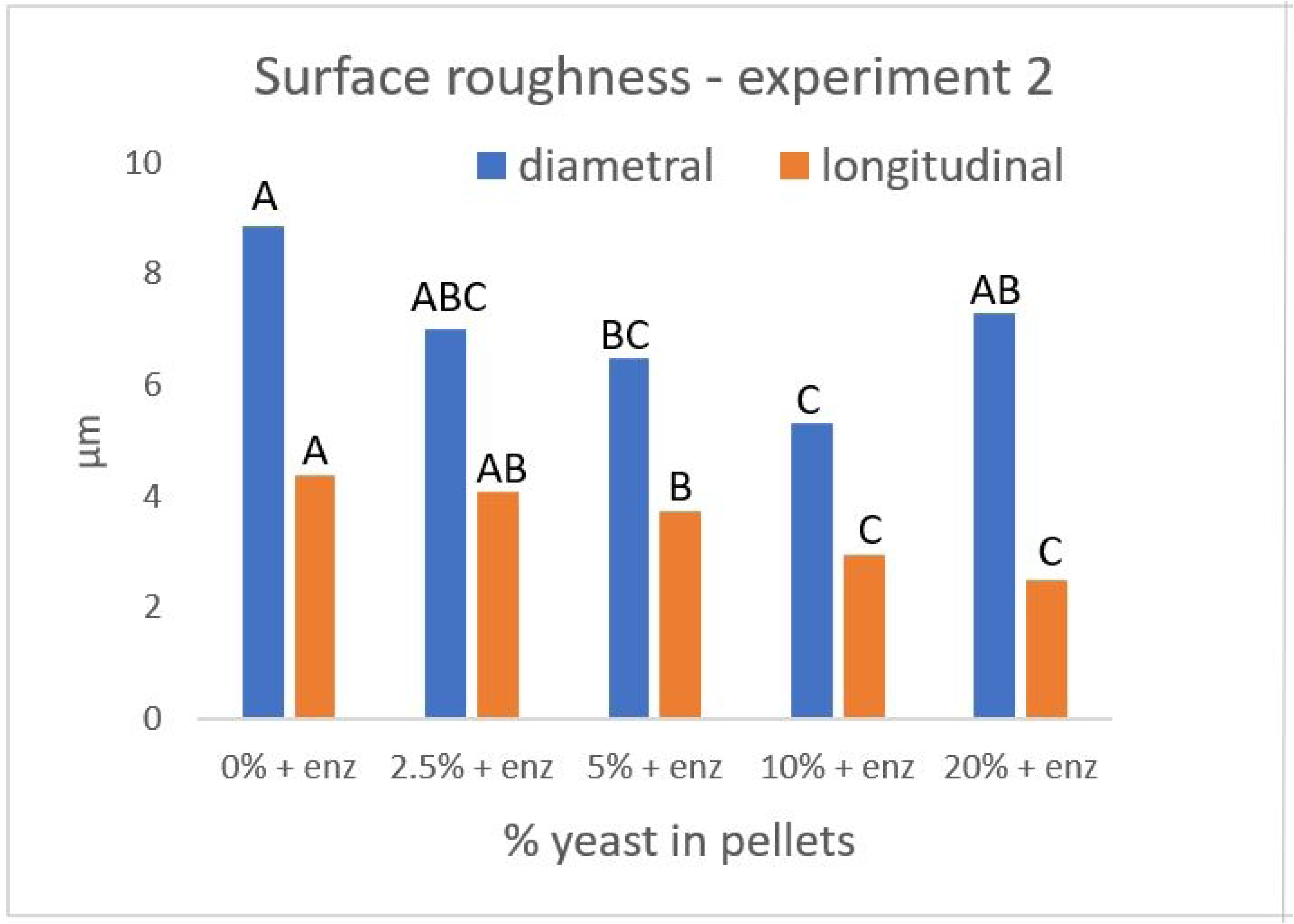
Surface roughness analyses data representing means based on 25 repetitions each for diametral and longitudinal analyses. Different sequential letters from ANOVA-Tukey statistical method signify difference at 5% level separately for diametral and for longitudinal measurements where p-values were lower than 0.001.

The surface roughness of pellets added enzymes, was not influenced p_max_, neither longitudinally (R^2^ = 0.08) nor diametrically (R^2^ = 0.01).

## 5. Discussion

Flow resistance (p_max_) in the die during pellet discharge in experiment 1 was shown to be independent of the addition from 2.5% up to 20% of the yeast *Cyberlindnera jadinii* (Table 2). When the yeast was the only medium to be pelleted, p_max_ showed to be significantly higher. This may be explained by changing particle size distribution of powder systems affecting their densification (relationship between particles) and compaction (relationship between particles and compaction die). When only yeast was pelleted the deposited micro-scale material of the single-cell organisms could possibly diffuse from one particle to another through solid bridges. Thus, the forces influencing the cohesion of powder particles will rest on the contact area of the solid bridges and their diameter, which could all influence the build-up of a pellet with a high-density structure (Pietsch, 1991). A large area of a compressed material may require high forces to be pushed through the channel of the die. A similar significant difference was observed in experiment 2 when the feed with 20% yeast was enzymatically treated (Table 3). A clear correlation between p_max_ and tensile strength was observed in experiment 2 (Fig. 7).

In experiment 1, adding up to 20% yeast to the feed significantly increased pellet tensile strength (Table 2). In experiment 2, enzymatic hydrolysis with a limited amount of water and 20% yeast generally enhanced the hardness of the feed pellets (Table 3). The observed improvement can be attributed to the intense interactions and compact packing of the smallest particles during the pelleting process, aligning with the findings of Miladinovic *et al*. (2013). Enhancing the durability of the pellets is likely to happen, among other factors, through interlocking bonds between the particles at the micro-scale.

Measurements done by surface contact angle analyses (CA) provided insight into how the structure of a compacted shrimp feed may absorb water or other liquids. CA measurements for oil and water showed to be partially dependent on the time of observation up to 94 seconds in experiment 1. A significant decrease of the oil CA was observed when 10% yeast was included (Table 2). A similar effect was observed for water after 47 seconds of observation. After 94 seconds even 2.5% added, yeast may contribute to significantly different CA for water. Comparable results were found for the CA of oil in experiment 2 after 10 seconds of observation. In experiment 2 the enzymatic hydrolysis and yeast inclusion decreased the CA for oil from 10 seconds and beyond 20 seconds of observation time, whereas the water CA showed very unclear results (Table 3). The reduction was observed only for 5% added yeast at 0.5 to 2 seconds of observation time. The entire set of results may be well explained by the density distribution of the initial powder and packing of the powder particles in the die (Burch *et al*., 2008). Also, Korachkin and Gethin (2004) explained that different packing of the particles may create two regions in compacted solids with densities differing by 10% during the die-fill. The authors observed such variation for various vertical die-fill density distributions with the same total mass filled in the die-hole. Such different final densities may also contribute to differing CA for both oil and water. The level of physical interaction between the feed pellets and different liquids in the presented research appears to depend on the friction, adhesion, adsorption, and wettability of the surface. This agrees with Yuan and Lee (2013).

Structural differences in the feed pellets caused by yeast and enzymes altered the properties of the pellet surface. Such alteration could be described by the diffusion between particles and liquid absorption in the cavities found between particles (Roman-Gutirrez *et al*., 2003). Defining disintegration of the feed pellets underwater is shown to be a good quality indicator of the usability of feed pellets by the shrimps, which agrees with Flemming (1995); Obaldo *et al*. (2002); and Bansemer *et al*. (2015). In the presented experimental work, it is shown that measuring the swelling rate through image analysis of undisturbed shrimp feed pellets under stagnant water developed by Salas-Bringas *et al*. (2015) and applied by Miladinovic *et al*. (2021) is a good tool. These measurements indicate the optimal time that a pellet can remain in the compact form before a shrimp eats it or before it disintegrates into the water.

The rate of underwater pellet swelling (UPS) is influenced by the adsorption and desorption of fluids, which are dependent on the physical and chemical properties of the compacted solids and the raw materials used in the pellets that form their surfaces. This is well described by Saalah *et al*. (2010). Chemical binding can either enhance or inhibit surface adsorption, with adsorption and desorption being continuous processes where molecules attach to and detach from surfaces. The observed large differences (37.3%) in UPS measurement for the first 60 seconds in the control treatments in both experiments with no yeast and without enzymatic treatment can be attributed to a 2% moisture difference. Determining the relationship between the rates of adsorption and desorption of molecules as influenced by external macroscopic variables, such as water vapor and its pressure, substrate temperature, and substrate surface structure was described by Oura *et al*. (2003). Free water in the pelleted matrix tends to accept faster the net movement of water molecules into pellets through a porous pellet structure. Molecular adsorption in compacted feed solids is the leading cause of swelling of hygroscopic materials such as non-starch polysaccharides. This is also described by Li *et al*. (2023), where the authors explain how mechanical stress affect the swelling behavior of hygroscopic materials. Non-starch polysaccharides significantly impact starch behavior, improving texture and functional properties by altering hydrolysis processes. The molecular adsorption process comes through the mechanisms of bound water, whereas capillary adsorption corresponds to free water. The free water molecules are drawn into the network of the walls of the compacted pellet structure by the forces of attraction. If these forces are large enough, they will draw the water molecule into the compacted structure. When the moisture of the compacted solids increases, the molecular attraction reduces, and therefore the volume increases. Such volume increase is closely equal to the volume of water added and hence the swelling of the pellets occurs faster. Water equilibrium is reached when molecules depart from the surface to the core of the pellets. Such equilibrium was achieved in experiment 1 after 20 minutes (Table 2). Enzymatic treatments influenced equilibrium after 40 minutes in experiment 2. There was no indication in experiment 1 that the replacement of fish meal with yeast influenced UPS. In experiment 2, a feed without yeast resulted in pellets being more prompt to swell underwater in the first 60 seconds (Table 3). Enzymatic treatment for 10% and 20% yeast inclusion had 48.6% lower UPS at the first minute, compared to feed without yeast. Similar results were found after 20 and 40 minutes. In the first experiment, the volume increase was not related to adding yeast as a raw material. However, by adding enzymes the shrimp feed with a higher content of yeast significantly slows the deterioration of compacted solids. The most likely explanation is that enzymes facilitated the formation of yeast microparticles into solids with minimal small cavities. These fissures could enhance water transport within the pellet through capillary adsorption, thereby altering the pellets water absorption capacity. Therefore, the pellet tensile strength increased over 3 folds by adding 20% yeast (Table 3).

In the longitudinal direction, the surface roughness differed only in the pellets containing 20% yeast in experiment 1. This may be attributed to a greater number of smallest particles that were able to pack better into a pellet. That is also following the increase of the tensile strength (Table 1). Enzyme addition contributed to the longitudinal decrease of roughness, which is observed for feed containing 5%, 10%, and 20% yeast (Fig. 10). Pellet surface roughness in the longitudinal direction was shown to be influenced by the particle size of the material, which is in line with Sarkar *et al*. (2014) and correlated with p_max_ during pelleting (Fig. 6).

## 6. Conclusions

Incorporating enzymatically untreated yeast at concentrations ranging from 2.5% to 20% in aquatic feed diets does not significantly affect the maximum pressure (p_max_) during pelleting, and thus, it is an indication that is very unlikely to increase electrical energy consumption compared to feed pellets containing fishmeal. However, adding 20% yeast is likely to produce harder pellets, which may extend the consumption time in water due to their slower swelling rate. This characteristic is beneficial for farms with slow-feeding aquatic organisms. To produce pellets with smoother surfaces, it is recommended to include 20% yeast in the feed diets. Enzymatic treatment of yeast increases p_max_ in feed diets containing 10% and 20% yeast, potentially leading to higher electrical energy usage. This treatment also enhances the hardness of pellets with 20% yeast and reduces their oil absorption capacity, attributed to the increased pellet hardness. The decreased longitudinal surface roughness of the pellet wall, achieved by replacing fishmeal with enzymatically treated yeast, may reduce the pellet’s ability to accept oil coating. The technical quality characterization of novel feed ingredients integrated into aquatic feeds may positively impact the physical quality of the pellets. This is crucial when dynamically changing feed diet formulations with varying inclusion levels of novel ingredients. To avoid high p_max_ in the die during feed pelleting, enzymatic hydrolysis should be avoided in feeds with 10% and 20% yeast inclusion. However, to increase the tensile strength of aquatic feed pellets, it is recommended to enzymatically treat yeast and include 20% in the formulation. Additionally, to decrease pellet swelling underwater between 20 to 40 minutes, feed pellets containing enzymatically treated yeast at 10% and 20% inclusion should be considered.

## Notes

### Competing Interest Statement

The authors have declared that no competing interests exist.

